# Palmitoylated importin α recruits PKCε to the plasma membrane to drive breast cancer cell motility

**DOI:** 10.64898/2026.05.28.728515

**Authors:** M. Kathryn Malone, Christopher W. Brownlee

**Affiliations:** Department of Pharmacological Sciences, Stony Brook University; Stony Brook, 11794, United States of America

## Abstract

Importin ⍺ is a nuclear transport factor which canonically has a role in binding and shuttling NLS-containing proteins from the cytoplasm into the nucleus. Recently, it has been shown that when palmitoylated by specific palmitoyl acyl transferases, importin ⍺ can partition to the plasma membrane where its roles remain widely unknown. Patients with breast cancer displaying increased importin ⍺ expression have advanced tumor size, poor tumor differentiation, and reduced overall and recurrence-free survival. In this study, we use palmitoylation altering pharmacological agents to demonstrate that membrane bound palmitoylated importin ⍺ enhances breast cancer cell motility through binding and tethering the serine/threonine kinase PKCε to the plasma membrane.

## INTRODUCTION

Breast cancer has the highest incidence rate of all forms of cancer globally and, in females, has the highest mortality rate globally (Bray et al., 2024; Siegel et al., 2026; Wagle et al., 2025). In the United States, it is the leading cause of cancer related death in Black and Hispanic women and is the second leading cause of cancer related death in women overall(Giaquinto et al., 2024; Siegel et al., 2026; Wagle et al., 2025). With increased early detection and the emergence of novel therapies targeting primary disease, the mortality rate for those with nonmetastatic disease continues to decrease each year (Bray et al., 2024; Giaquinto et al., 2024; Siegel et al., 2026; Wagle et al., 2025). Despite this progress, patients with distant metastases have just 21-31% 5-year survival rate indicating a large demand for metastatic progression and metastasis-targeting therapies (Burcu et al., 2023; Giaquinto et al., 2024; Siegel et al., 2026; Wagle et al., 2025; R. Wang et al., 2019).

Recently, the membrane bound *O*-acyltransferase (MBOAT) protein porcupine (PORCN) has been the target of novel emerging cancer therapies (Black et al., 2025; Boone et al., 2016; Lanyon-Hogg et al., 2017a; Li et al., 2020; D. Liu et al., 2019; Madan et al., 2015; Neiheisel et al., 2022; Rodon et al., 2021; Serafino et al., 2017; Shah et al., 2021). PORCN is responsible for the palmitoylation, or reversible post-translational addition of a palmitate fatty acid on the serine residues of its primary target protein, Wnt, as well as its newly discovered target, importin ⍺ (Brownlee & Heald, 2019; Y. Liu et al., 2022). The family of 24 zDHHC lipid transferases, which are responsible for cysteine S-palmitoylation, has also been implicated in their involvement in cancer progression (Lanyon-Hogg et al., 2017b; Lin, Agarwal, et al., 2023; Tang et al., 2022; Tomić et al., 2024; Yeste-Velasco et al., 2014). Despite these emerging findings, there still remain large gaps in understanding the complete involvement of these enzymes in cancer progression (Ko & Dixon, 2018).

The nuclear import factor importin ⍺ is a member of the karyopherin (KPN) transport family of proteins, which is comprised of seven human importin ⍺ subtypes (KPNA1-KPNA7) and two importin β subtypes (KPNB1 and KPNB2) (Miyamoto et al., 2016; Oka & Yoneda, 2018; Pumroy & Cingolani, 2015). The primary role of importin ⍺ is to transport proteins into the nucleus by binding importin β then forming a ternary complex with nuclear localization signal (NLS)-containing proteins in the cytosol and subsequently translocating into the nucleus through the nuclear pore complex. Once in the nucleus, the proteins dissociate, the cargo is released, and importin ⍺ and importin β are recycled to the cytosol (Miyamoto et al., 2016; Oka & Yoneda, 2018). Recently, advancements have been made in understanding alternative functions for one member of the importin ⍺ family in particular, KPNA2 (Oka & Yoneda, 2018; Pumroy & Cingolani, 2015). Notably, the involvement of KPNA2 (importin ⍺) has been indicated in various forms of cancer and its upregulation has been implicated as a biomarker for poor prognoses (Alshareeda et al., 2015; Christiansen & Dyrskjøt, 2013; Han et al., 2021; Noetzel et al., 2012; Pumroy & Cingolani, 2015; C. I. Wang et al., 2011; Yamada et al., 2016). Additionally, overexpression of KPNA2 in breast cancer cells *in vitro* resulted in a significant increase in both proliferation and migration. Notably, overexpression in benign breast cells *in vitro* not only caused a significant increase in these metrics but also resulted in reduced cell-matrix adhesion. These findings collectively contribute to increased colony spreading, thereby indicating a shift towards a malignant phenotype (Noetzel et al., 2012).

KPNA2 has also been identified at the cell surface in several cancer types where it has been observed to retain its ability to bind NLS containing cargo, including certain growth factors, at the plasma membrane (PM) (Oka & Yoneda, 2018; Olsnes et al., 2003; Yamada et al., 2016). These findings suggest one potential mechanism for promoting cancer progression. More recently, KPNA2 has been shown to localize to the PM via palmitoylation, where it has been shown to play roles in organelle size control, ciliogenesis, and mitotic spindle orientation (Brownlee & Heald, 2019; Mosqueda et al., 2025; Oka & Yoneda, 2018; Olsnes et al., 2003; Sutton et al., 2025; Yamada et al., 2016). These studies have identified four palmitoylation sites within KPNA2, two serine and two cysteine, that are conserved in *Xenopus laevis* and humans. PORCN is responsible for the serine palmitoylation of KPNA2, while the cysteine palmitoylation is performed by one or more unidentified zDHHC transferases (Brownlee & Heald, 2019; Sutton et al., 2025). Given the pro-tumorigenic effects of PORCN and various zDHHC transferases, they have been investigated as potential targets for cancer treatment (Boone et al., 2016; Li et al., 2020; Lin, Agarwal, et al., 2023; Lin, Lv, et al., 2023; D. Liu et al., 2019; Y. Liu et al., 2022; Madan et al., 2015; Neiheisel et al., 2022; Proffitt et al., 2013; Rodon et al., 2021; Serafino et al., 2017; Shah et al., 2021; Tang et al., 2022; Tomić et al., 2024; Yeste-Velasco et al., 2014). Additionally, the drugs Wnt-C59 and ivermectin have both been implicated as anti-cancer treatments with both drugs affecting importin ⍺ through inhibition of PORCN mediated palmitoylation and importin ⍺ cargo binding, respectively (Dutra et al., 2023; Jiang et al., 2019; Juarez et al., 2020; Lee et al., 2016; Shah et al., 2021; Wagstaff et al., 2011, 2012).

Importin ⍺ serves as a connecting factor in various emerging therapies. Therefore, in this work, we posit and demonstrate that palmitoylated importin ⍺ regulates metastasis through its interactions at the PM. We show that palmitoylated importin ⍺ is required at the PM for cell motility. We also identify a novel interaction between importin ⍺ and the predicted NLS region within the serine/threonine kinase protein kinase C ε (PKCε), which has been previously linked to alterations in cell motility and metastatic disease (Gorin & Pan, 2009; Jansen et al., 2001; Mischaks et al., 1993). We further find this interaction to be vital for membrane localization of PKCε. Additionally, we show membrane localized PKCε to modulate motility through a mechanism preferentially used by mesenchymal cells over benign cells. Together, this work elucidates a previously unexplored mechanism for mesenchymal cell motility that holds potential for future drug targets.

## RESULTS

### Altering importin ⍺ palmitoylation status and function affects cell motility and invasion

Overexpressed importin ⍺ has been identified as a marker of poor prognosis in human breast cancer and is associated with increased rates of metastasis (Alshareeda et al., 2015; Christiansen & Dyrskjøt, 2013; Han et al., 2021; Noetzel et al., 2012; Pumroy & Cingolani, 2015; C. I. Wang et al., 2011; Yamada et al., 2016). When importin ⍺ is overexpressed in both benign and malignant breast cell lines, it promotes cell motility and reduces cell-matrix adhesion (Alnoumas et al., 2021; Christiansen & Dyrskjøt, 2013; Noetzel et al., 2012). Notably, PORCN is responsible for the serine palmitoylation of importin ⍺, which directs it to the PM where it has been identified in several cancer types and maintains its ability to bind NLS containing proteins (Brownlee & Heald, 2019; Oka & Yoneda, 2018; Olsnes et al., 2003; Yamada et al., 2016). PORCN has also been investigated as a potential target for cancer treatment (Boone et al., 2016; Li et al., 2020; Madan et al., 2015; Proffitt et al., 2013; Rodon et al., 2021; Shah et al., 2021). Therefore, we sought to investigate the potential role of palmitoylation of importin ⍺ by PORCN in breast cancer metastasis. To probe the mechanism by which importin ⍺ may affect the metastatic qualities of breast cancer cells, a panel of drugs was utilized to separately alter importin ⍺ palmitoylation status and function.

Importin ⍺ is one of two known targets of O-palmitoylation by the MBOAT PORCN, making PORCN a logical inhibitory target to decrease the palmitoylated, membrane bound fraction of importin ⍺ (Proffitt et al., 2013). To accomplish this, a small molecule inhibitor, Wnt-C59, was used to specifically target PORCN and inhibit O-palmitoylation of importin ⍺. Additionally, the small molecule inhibitor of protein acyl transferases (PATs), 2-bromopalmitate, was used to selectively inhibit broad spectrum s-palmitoylation and thus lower the palmitoylated fraction of all proteins (Davda et al., 2013). To increase the palmitoylated fraction of importin ⍺, palmostatin B was used, which inhibits the acyl protein thioesterase APT1, responsible for depalmitoylating importin ⍺ (Dekker et al., 2010). Finally, importin ⍺ function was impaired with the small molecule inhibitor importazole which prevents nuclear import of NLS-containing proteins by disrupting interactions with RanGTP (Soderholm et al., 2011). Treatment with Wnt-C59 and importazole both significantly decreased cell motility as analyzed using a wound healing assay (**Fig. 1A,B**). Wnt-C59 treatment directly inhibits importin ⍺ from localizing to the plasma membrane; it is plausible that importazole treatment indirectly inhibits importin ⍺ localization to the PM by inhibiting the importin ⍺/β-cargo complex from dissociating within the nucleus thereby reducing the fraction of cytosolic importin ⍺ that is free to be palmitoylated and sent to the PM. Additionally, altering the palmitoylation status of importin ⍺ either to decrease the palmitoylated fraction with Wnt-C59 or globally decreasing protein palmitoylation via 2-bromopalmitate both reduce the ability of cells to invade through a Matrigel coated membrane (**Fig. 1C,D**). This is indicative of a required balance of palmitoylated membrane bound to depalmitoylated cytosolic importin ⍺ for the maintenance of normal cancer cell invasion. To ensure the drug-induced changes seen in motility and invasion were not a byproduct of changes in cell cycle, cells were stained for the G2-M progression marker phospho-Histone H3 (pHH3) and the ratio of pHH3 positive cells to total cells counted was calculated (**Fig. 1E**). There were no observed changes in cell cycle upon drug treatment. These data demonstrate the ability to inhibit chemoattractant or wound healing-related directional cell motility.

**Figure 1.**
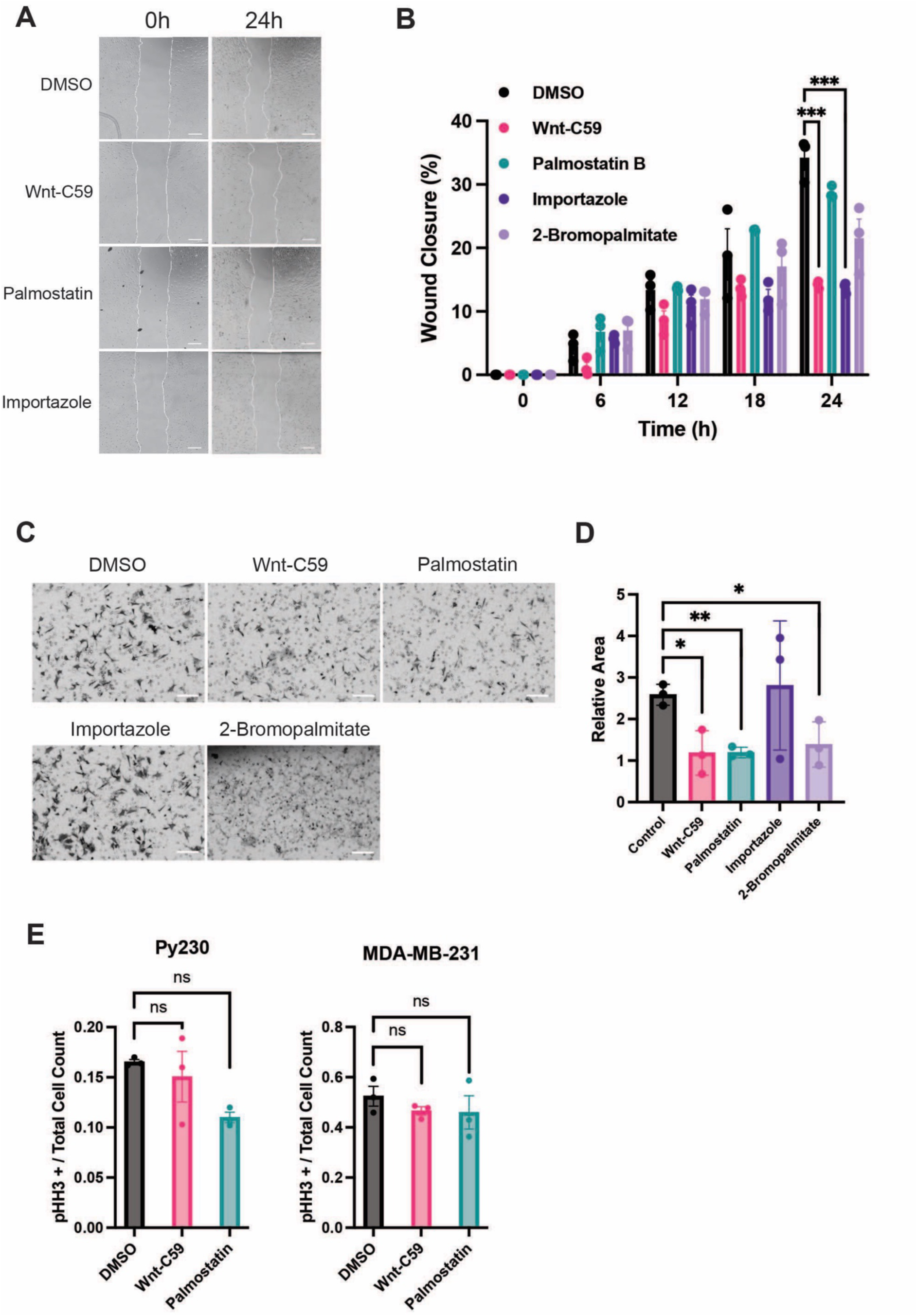
Altering importin ⍺ palmitoylation status and function affects cell motility and invasion. **A)** Phase-contrast images of Py230 cells at time 0h and 24h of migration assay following treatment with DMSO, 10µM Wnt-C59, 50µM palmostatin, 40µM importazole, or 100µM 2-bromopalmitate for 24 hours prior to analysis with the wound border marked in white. Scale bars = 500µm. **B)** Quantification of wound closure in migration assay with Py230 cells. Images taken every 6 hours for 24 hours. ***P<0.001, Student’s t-test, 3 replicates per condition. Mean ± SEM. **C)** Brightfield images of Py230 cells treated with DMSO, 10µM Wnt-C59, 50µM palmostatin, 40µM importazole, or 100µM 2-bromopalmitate for 24 hours prior to analysis in transwell invasion assay. Cells pictured successfully migrated through the Matrigel coated transwell membrane and were stained with crystal violet. Scale bars = 100µm. **D)** Quantification of invasive cells in transwell invasion assay. *P<0.05, **P<0.01, Student’s t-test, 3 replicates per condition. Mean ± SEM. **E)** Quantification of the ratio of pHH3 positive cells to total cells counted in either Py230 or MDA-MB-231 cells following treatment with DMSO, 10µM Wnt-C59, or 50µM palmostatin for 24 hours. One-way ANOVA with Dunnett’s multiple comparisons test, 3 replicates per condition, 100 cells per replicate. Mean ± SEM.

### Palmitoylated importin ⍺ is required for cancer cell motility

Inhibition of importin ⍺ palmitoylation resulted in diminished wound healing and invasion thus demonstrating the migratory ability of whole cell masses and the ability of cells to recognize and respond to chemoattractants and properly adhere to extracellular matrix. Despite these findings, it remains unclear which specific processes are defective upon importin ⍺ palmitoylation inhibition. To address this and determine the effects on motility independent of directionality, using the cells’ ability to detect nutrients, and direct cell-cell communication, a 2D individual cell tracking approach was used via live cell imaging. In this study, cells were individually tracked over a period of 20 hours (**Fig. 2A**). Following treatment with Wnt-C59, Py-230 cells had a dramatically lower average accumulated distance and velocity (**Fig. 2B,C**). This demonstrates a more direct role for palmitoylated importin ⍺ in individual cell motility in general.

**Figure 2.**
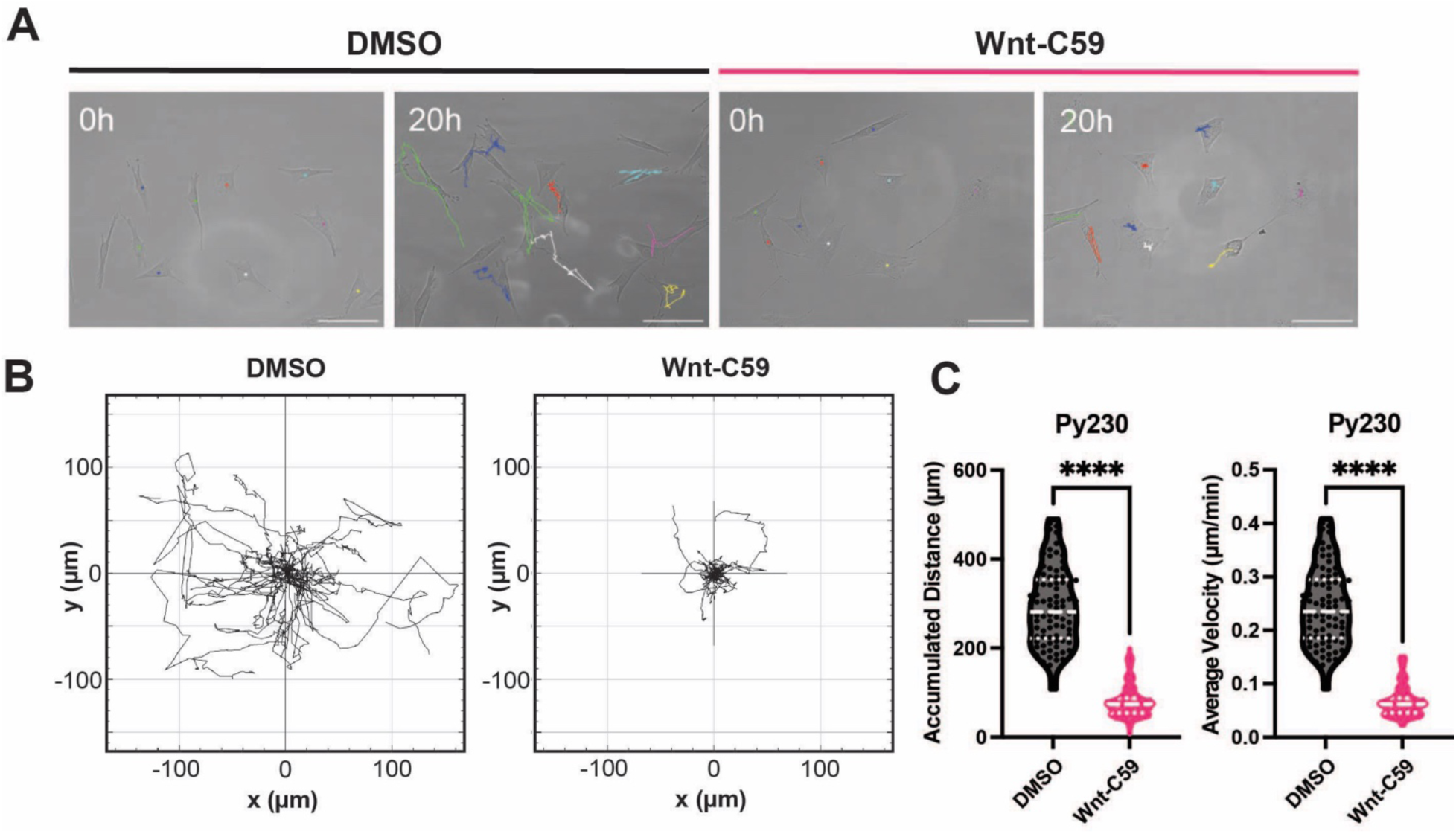
Palmitoylated importin ⍺ is required for normal cancer cell motility. **A)** Phase-contrast images of Py230 cells treated with DMSO or 10µM Wnt-C59 at initial (0h) and end (20h) time with overlaid trajectories of 9 cells. Images captured with time-lapse microscopy at 20-minute intervals. Scale bars = 100µm. **B)** Trajectory plots showing Py230 cell trajectories over 20h in DMSO and Wnt-C59 treated cells. Cell tracks were all set to a common origin. **C)** Quantification of accumulated distance (µm) and average velocity (µm/h) of Py230 cells treated with DMSO or Wnt-C59 over 20h. ****P<0.0001, Student’s t-test, n=90 cells, 3 replicates per condition. Median and quartiles marked by dashed lines.

To further elucidate if the observed changes in cell motility were due to the restriction of importin ⍺ from localizing to the membrane following Wnt-C59 treatment and not off-target effects, importin ⍺-HA-mCherry (importin ⍺-WT) and importin ⍺-HA-mCherry-CaaX (importin ⍺-CaaX) constructs were utilized. The CaaX motif is comprised of a cysteine residue, two aliphatic residues, and a C-terminal amino acid. This motif is present on the C-terminus of a group of proteins that are involved in various regulatory and signaling processes in which membrane localization is critical. The CaaX motif marks proteins for prenylation, specifically farnesylation, which subsequently targets the proteins to the PM where they then carry out their respective functions (Gao et. al.). The addition of the CaaX motif to importin ⍺ allows for its targeting to the PM independent of palmitoylation status. MDA-MB-231 cells were transfected with either the importin ⍺-WT or importin ⍺-CaaX plasmid and were treated with DMSO or Wnt-C59. The cells transfected with importin ⍺-WT behaved as previously observed following drug treatment in that administration of Wnt-C59 caused a decrease in migratory distance and velocity compared to those of DMSO treated. There were no differences in either parameter between cells transfected with importin ⍺-WT and importin ⍺-CaaX when treated with DMSO. Interestingly, cells transfected with importin ⍺-CaaX and treated with Wnt-C59 displayed a hyper-motile phenotype compared to that of all other conditions (**Fig. 3A-C**). This demonstrates the ability of membrane-bound importin ⍺ to rescue the reduced cell motility seen when palmitoylation is inhibited. Importantly, this suggests that the inhibition of motility upon PORCN inhibition is due solely to preventing palmitoylation of importin ⍺.

**Figure 3.**
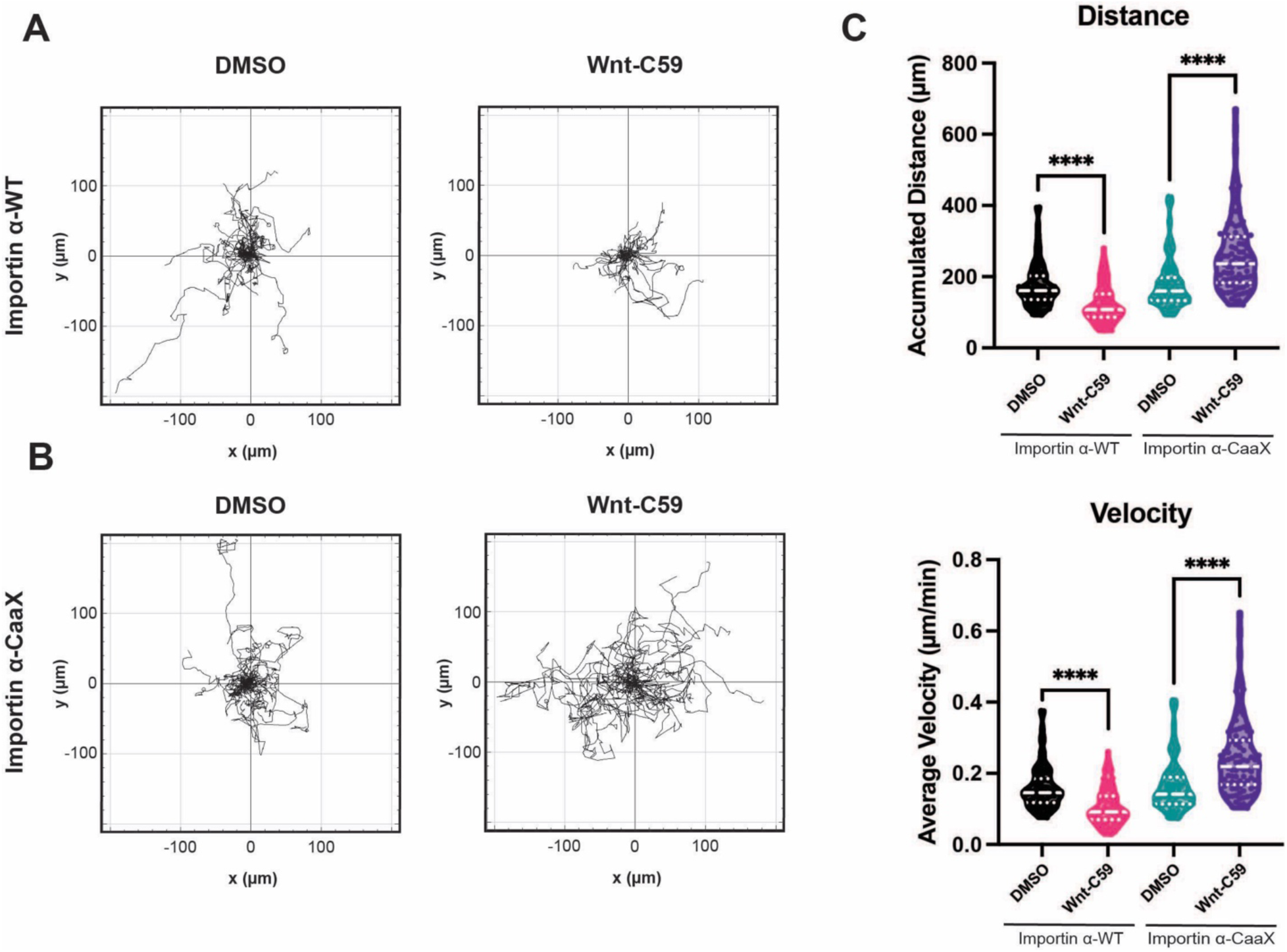
Forcing importin ⍺ to the PM reverses the non-motile phenotype seen with Wnt-C59 treatment. **A)** Trajectory plots showing the trajectories of MDA-MB-231 cells transfected with importin ⍺-WT or **B)** importin ⍺-CaaX and treated with DMSO or 10µM Wnt-C59 over 20 hours. Cell tracks were all set to a common origin. **C)** Quantification of accumulated distance (µm) and average velocity (µm/h) of transfected MDA-MB-231 cells treated with DMSO or Wnt-C59 over 20h. ****P<0.0001, One-way ANOVA with Tukey’s multiple comparisons test, n=90 cells, 3 replicates per condition. Median and quartiles marked by dashed lines.

### NLS prediction scores of proteins identified at the PM that impair cell migration upon siRNA knockdown

In its role as a nuclear import factor, importin ⍺ complexes with importin β to allow for binding of cytoplasmic proteins which contain NLS prior to translocation into the nucleus where the proteins are then released from the complex. In a previous study, a genome-wide screen for NLS-containing proteins identified an enriched population at the PM (Sutton et al., 2025). This evidence, along with the knowledge that importin ⍺ is present at the PM in various cancers while maintaining its ability to bind NLS cargo (Oka & Yoneda, 2018; Olsnes et al., 2003; Yamada et al., 2016), raises the intriguing hypothesis that importin ⍺ binds NLS-containing cargo in the cytoplasm and helps to tether the cargo to the PM upon palmitoylation. To identify potential candidates for importin ⍺ binding, a list of previously identified (Simpson et al., 2008) genes that impair cell migration when knocked down with siRNA was further screened for localization to the PM. Of these, PKCε was identified to likely contain a NLS (**Table 1**). PKCε is a member of the protein kinase C (PKC) family of serine/threonine kinases. Notably, PKCε has been previously linked to alterations in cell motility and metastatic disease. When PKCε is overexpressed in immortalized epithelial cells, there are marked increases in motility and invasion(Mischaks et al., 1993) and overexpression in animal models yields increased tumorigenic and metastatic phenotypes(Jansen et al., 2001). PKCε is also overexpressed in tumor-derived cell lines and histopathological tumor specimens (Gorin & Pan, 2009).

**Table 1.**
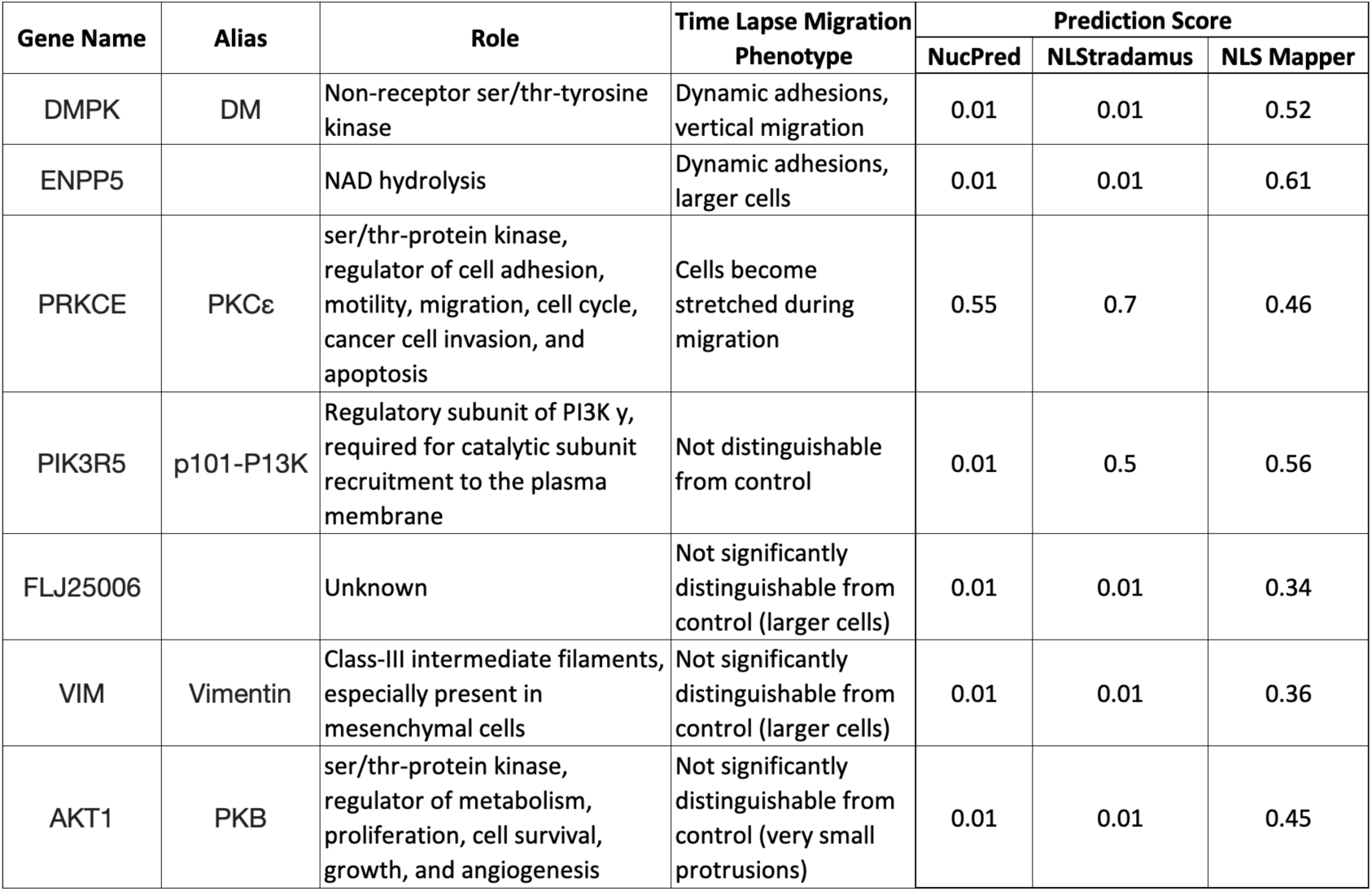
NLS prediction scores of proteins identified at the PM that impair cell migration upon siRNA knockdown.

### PKCε PM localization requires NLS-mediated binding to palmitoylated importin ⍺

PKCε must first be transported to the PM where it is activated by diacyl glycerol (DAG) however, the exact mechanism through which it is localized is unknown. In order to explore the possibility of its PM localization being regulated by palmitoylated importin ⍺, we first sought to determine if importin ⍺ could bind to PKCε. Through immunoprecipitation of transfected PKCε-HA and endogenous importin ⍺, we found that importin ⍺ is able to immunoprecipitate with PKCε (**Fig. 4B**). To confirm that PKCε requires its predicted NLS to immunoprecipitate with importin ⍺, we generated a mutant PKCε construct in which glycine and alanine were substituted with the charged arginine and lysine residues within the predicted NLS motif (**Fig. 4A**). A ubiquitination site exists within the predicted NLS sequence and so we did not mutate the lysine to avoid potential adverse effects on protein stability or function. Immunoprecipitation of the transfected mutant PKCε-HA-ΔNLS and endogenous importin ⍺ showed that the two proteins could no longer be co-immunoprecipitated, thus confirming PKCε requires its predicted NLS sequence to be immunoprecipitated with importin ⍺ (**Fig. 4C**). To determine if PKCε localization to the PM is affected by importin ⍺ palmitoylation, we treated cells transfected with PKCε-HA with the palmitoylation altering drugs Wnt-C59 and palmostatin. When cells were treated with Wnt-C59, thus decreasing the palmitoylated, PM-bound fraction of importin ⍺, the ratio of PKCε-HA at the PM to cytoplasm was also decreased (**Fig. 4D,E**). To further confirm the role of importin ⍺ on PKCε PM localization, we repeated the experiment using cells transfected with PKCε-HA-ΔNLS. Treatment with Wnt-C59 in these cells showed no difference in PKCε-HA-ΔNLS PM localization from DMSO, suggesting that the predicted NLS is required for PKCε PM localization (**Fig. 4F, G**). Additionally, to confirm that localization of PKCε to the PM requires importin ⍺ palmitoylation, MDA-MB-231 cells were transfected with either importin ⍺-WT or importin ⍺-CaaX, treated with DMSO or Wnt-C59, and were fixed and stained for PKCε. Cells transfected with importin ⍺-WT displayed a similar significant decrease in PM to cytoplasmic ratio of PKCε following Wnt-C59 treatment when compared to cells only expressing endogenous importin ⍺ (**Fig. 4H,I**). Cells expressing importin ⍺-CaaX notably displayed the opposite phenotype when treated with Wnt-C59 in that the PM to cytoplasmic ratio of PKCε increased (**Fig. 4J,K**). Together, these data demonstrate the novel relationship between importin ⍺ and PKCε in that importin ⍺ binds to the NLS on PKCε and localizes it to the PM following palmitoylation.

**Figure 4.**
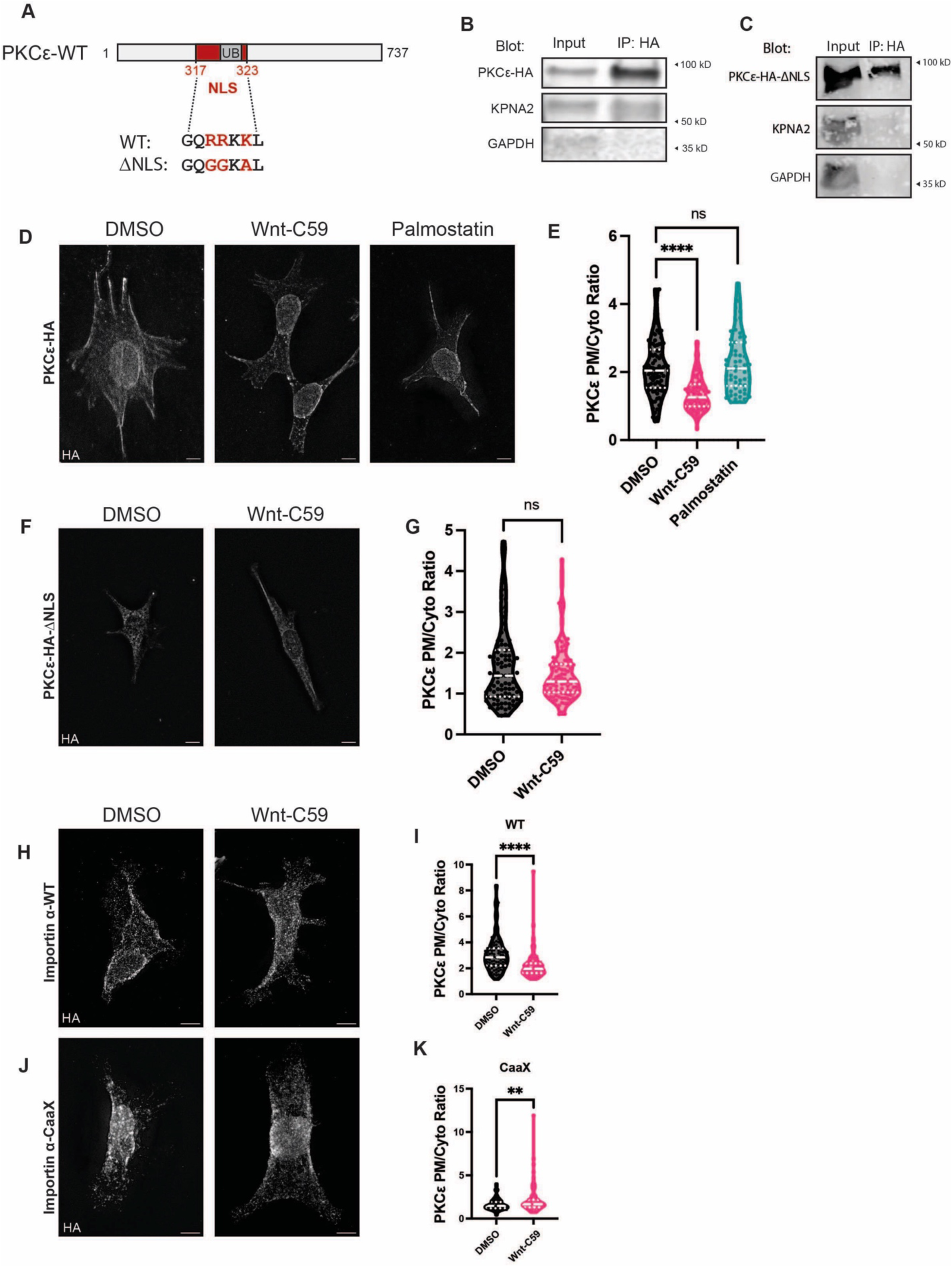
Importin ⍺ binds to PKCε via its NLS and localizes it to the PM when palmitoylated. **A)** Schematic overview of PKCε with the predicted NLS sequence spanning amino acids 317 to 323. PKCε-ΔNLS mutations made are shown in red while the ubiquitination site in gray was not mutated. **B)** Immunoblot of Py230 cells transfected with PKCε-HA and immunoprecipitated for HA followed by PKCε, KPNA2 (importin ⍺), and GAPDH western blot. **C)** Immunoblot of Py230 cells transfected with PKCε-HA-ΔNLS and immunoprecipitated for HA followed by PKCε, KPNA2, and GAPDH western blot. **D)** Fluorescence images of Py230 cells transfected with PKCε-HA and treated with DMSO, 10µM Wnt-C59, or 50µM palmostatin for 24 hours prior to immunostaining for HA. Scale bars = 10µm. **E)** Quantification of the ratio of PM to cytoplasmic localization of PKCε-HA in transfected Py230 cells. ****P<0.0001, one-way ANOVA with Dunnett’s multiple comparisons test, n=90 cells, 3 replicates per condition. Median and quartiles marked by dashed lines. **F)** Fluorescence images of Py230 cells transfected with PKCε-HA-ΔNLS and treated with DMSO or 10µM Wnt-C59 for 24 hours prior to immunostaining for HA. Scale bars = 10µm. **G)** Quantification of the ratio of PM to cytoplasmic localization of PKCε-HA-ΔNLS in transfected Py230 cells. Student’s t-test, n=90 cells, 3 replicates per condition. Median and quartiles marked by dashed lines. **H)** Fluorescence images of MDA-MB-231 cells transfected with importin ⍺-WT and treated with DMSO or 10µM Wnt-C59 for 24 hours prior to immunostaining for PKCε. Scale bars = 10µm. **I)** Quantification of the ratio of PM to cytoplasmic localization of PKCε in transfected MDA-MB-231 cells. ****P<0.0001, Student’s t-test, n=90 cells, 3 replicates per condition. Median and quartiles marked by dashed lines. **J)** Fluorescence images of MDA-MB-231 cells transfected with importin ⍺-CaaX and treated with DMSO or 10 µM Wnt-C59 for 24 hours prior to immunostaining for PKCε. Scale bars = 10µm. **K)** Quantification of the ratio of PM to cytoplasmic localization of PKCε in transfected MDA-MB-231 cells. **P<0.01, Student’s t-test, n=90 cells, 3 replicates per condition. Median and quartiles marked by dashed lines.

### Activating PKCε increases motility in untransformed but not cancerous breast epithelial cells

PKCε is activated by DAG at the PM thus enabling it to phosphorylate its local targets. To assess the effects of PKCε activity on motility, we utilized the selective PKCε agonist dicyclopropyl-linoleic acid (DCP-LA) and tracked the location of individual cells over the course of 20 hours. In the malignant breast cancer cell line, Py230, treatment with DCP-LA resulted in no change of the accumulated distance or average velocity compared to those of control treated cells. Additionally, there was no difference in accumulated distance or average velocity in cells treated with Wnt-C59 or the combination of Wnt-C59 and DCP-LA (**Fig. 5A,B**). Together, these results indicate that PKCε is maximally active in malignant Py230 cells. We then wanted to assess effects of PKCε activity in the benign breast cell line EpH4. In these cells, treatment with DCP-LA significantly increased both accumulated distance and average velocity compared to control. This increased motility was able to be attenuated with Wnt-C59 treatment. Notably, treatment with Wnt-C59 alone failed to diminish accumulated distance or average velocity when compared with control treatment (**Fig. 5C,D**). These results are indicative of separate migratory mechanisms between invasive, malignant breast cancer cells vs normal breast epithelial cells.

**Figure 5.**
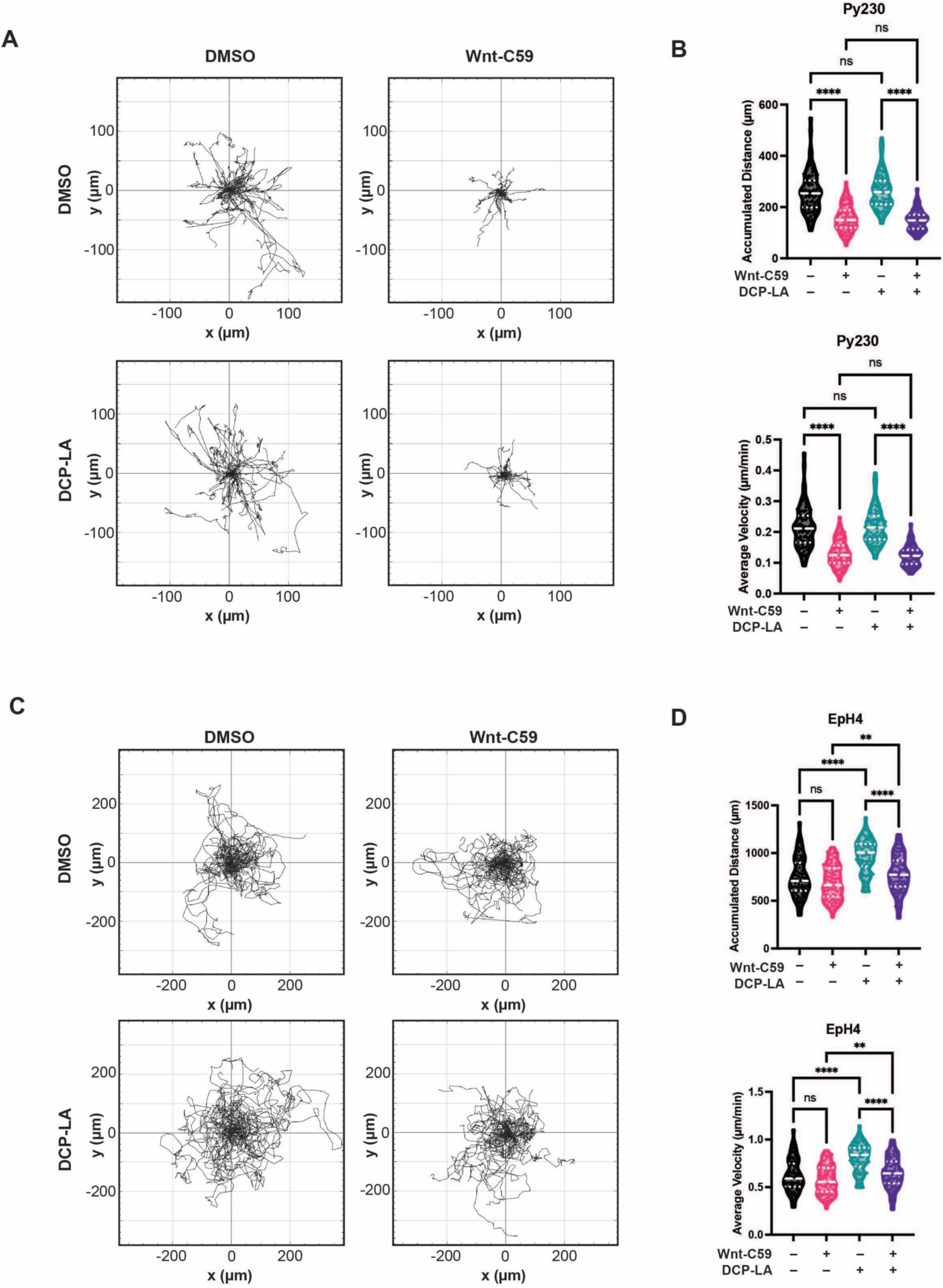
PKCε is inactive in normal epithelial cells and active in cancer cells as a driver of motility. **A)** Trajectory plots of Py230 cells treated in combination with DMSO, 10µM Wnt-C59, and/or 500nM DCP-LA. Plots represent individual cell locations and trajectories over 20 hours at 20-minute intervals. Cell tracks were set to a common origin. **B)** Quantification of accumulated distance (µm) and average velocity (µm/h) of Py230 cells treated with DMSO, Wnt-C59, and/or DCP-LA over 20h. ns=not significant, ****P<0.0001, one-way ANOVA with Tukey’s multiple comparisons test, n=90 cells, 3 replicates per condition. Median and quartiles marked by dashed lines. **C)** Trajectory plots of EpH4 cells treated in combination with DMSO, 10µM Wnt-C59, and/or 500nM DCP-LA. Plots represent individual cell locations and trajectories over 20 hours at 20-minute intervals. Cell tracks were set to a common origin. **D)** Quantification of accumulated distance (µm) and average velocity (µm/h) of EpH4 cells treated with DMSO, Wnt-C59, and/or DCP-LA over 20h. **P<0.01, ****P<0.0001, one-way ANOVA with Tukey’s multiple comparisons test, n=90 cells, 3 replicates per condition. Median and quartiles marked by dashed lines.

## DISCUSSION

Cancer therapies targeting PORCN have been developed with the assumptions that the anti-tumorigenic phenotypes observed are due solely to inhibition of Wnt secretion (Black et al., 2025; Boone et al., 2016; Lanyon-Hogg et al., 2017b; Li et al., 2020; D. Liu et al., 2019; Madan et al., 2015; Neiheisel et al., 2022; Rodon et al., 2021; Serafino et al., 2017; Shah et al., 2021). Our study identifies an additional pathway by which inhibition of PORCN has anti-tumor effects through inhibition of importin ⍺ palmitoylation. Importin ⍺ is classically recognized for its role as a nuclear import factor, binding NLS sequence-containing proteins and translocating them into the nucleus. Recent studies have described alternative processes involving importin ⍺ when palmitoylated and localizing to the PM(Brownlee & Heald, 2019; Mosqueda et al., 2025; Sutton et al., 2025). In the present study, we have demonstrated a novel mechanism through which importin ⍺ regulates breast cancer cell motility (**Fig. 6**). We have shown that disrupting importin ⍺ palmitoylation, and thus reducing its presence at the PM, diminishes PKCε membrane localization and activity resulting in decreased cell motility.

**Figure 6.**
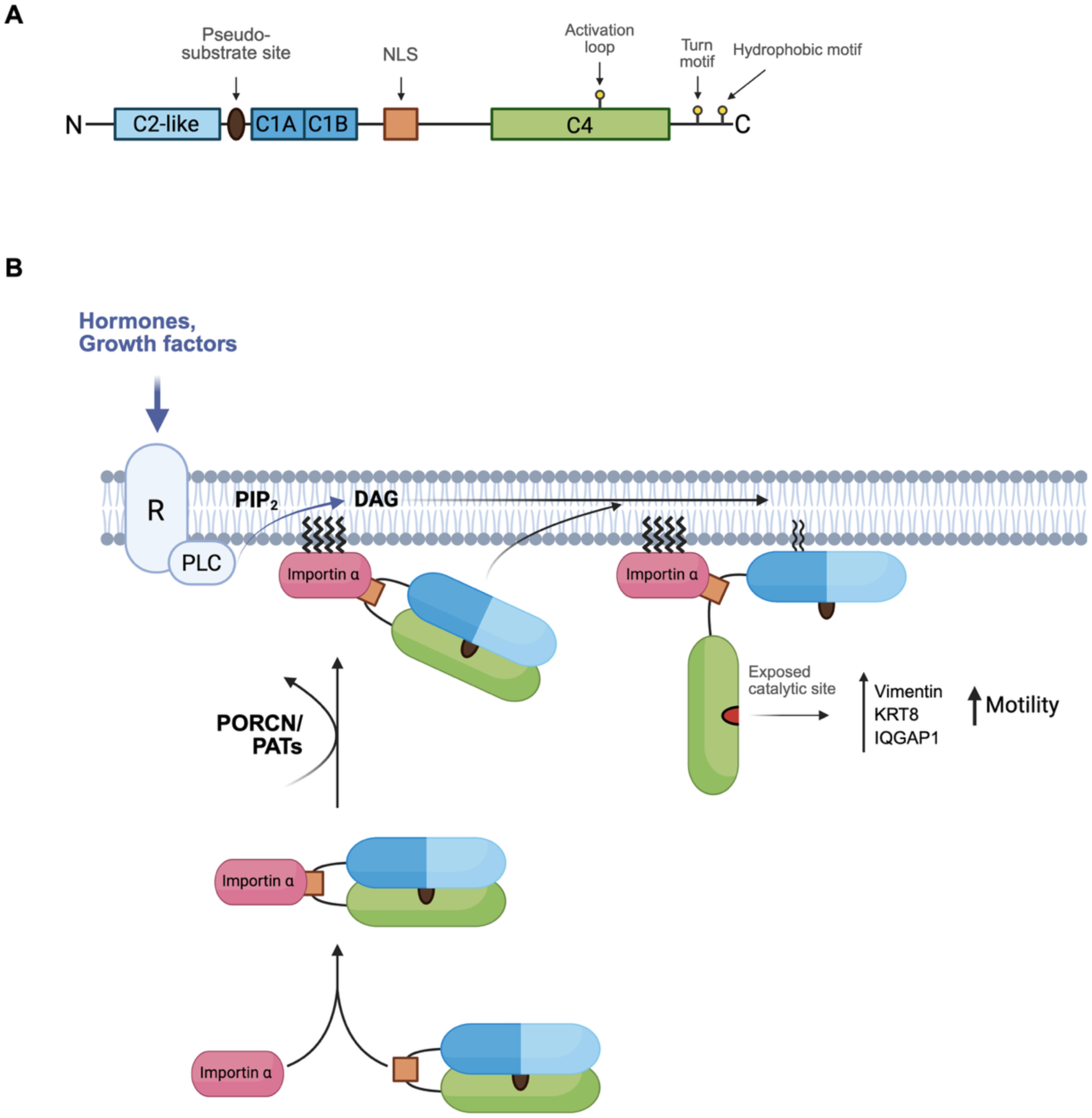
Importin ⍺ forms a complex with PKCε and tethers it to the PM upon palmitoylation of importin ⍺ allowing PKCε to become activated and increase breast cancer cell motility. **A)** Domain composition of PKCε showing the C2-like domain (light blue), the pseudo-substrate site (brown), the C1A and C1B domains (blue; responsible for binding DAG), the proposed NLS (orange), the C4 domain (green; kinase domain), and the three priming phosphorylation sites: the activation loop located in the C4 domain and the turn motif and hydrophobic motif located in the C-terminal tail (yellow circles). **B)** Proposed mechanism through which importin ⍺ tethers PKCε to the PM where it becomes catalytically active to phosphorylate downstream factors and increase breast cancer cell motility.

We first demonstrated that importin ⍺ is required at the membrane for individual and mass cell migration and invasion through the use of pharmacological agents (**Fig. 1, 2)**. By using CaaX modified importin ⍺ constructs, we showed that the effects on motility were not dependent on palmitoylation, rather on the ability of importin ⍺ to localize to the PM (**Fig. 3**). Importin ⍺ is involved in a number of cellular functions and processes through NLS-binding and subsequent regulation by localization or by acting as a competitive inhibitor through blocking binding sites overlapping a protein’s NLS sequence (Oka & Yoneda, 2018). Importin ⍺ has also been found to be present at the PM and maintain its ability to bind NLS-containing cargo in various cancers (Oka & Yoneda, 2018; Olsnes et al., 2003; Yamada et al., 2016). Thus, we hypothesized that importin ⍺ may be affecting cell migration and invasion by interacting with an NLS-containing protein already present at the PM or alternatively transporting such a protein to the PM where it then becomes active.

To identify potential binding partners, we screened a group of genes shown to impair cell motility upon siRNA knockdown (Simpson et al., 2008) for their localization to the PM. Those positive hits were further assessed for their predicted likelihood of containing an NLS sequence. Through this process, the serine/threonine kinase PKCε was predicted to contain an NLS sequence in its hinge region (Table 1). PKCε is present in the cytosol in an inactive conformation and is recruited to the PM where it is allosterically activated by DAG (Acs et al., 1997; Bogi et al., 1998; Chun et al., 1996; Dries et al., 2007; Giorgione et al., 2006; Johnson et al., 2000; Lu et al., 1998; Lučić et al., 2016; Newton, 2018; Schmitz-Peiffer, 2020). The mechanism by which it is recruited to the PM is not well understood. Conflicting reports exist on the topic; some studies suggest that PKCε is transported to the PM solely through its affinity to phosphatidylserine (PS) which is enhanced in the presence of DAG while other studies suggest that PKCε has generally low affinity and specificity for PS. All of these studies draw conclusions using vesicles containing PS at high molar ratios yet show minimal affinity at the lower physiological levels of PS (Giorgione et al., 2006; Johnson et al., 2000; Leventis & Grinstein, 2010; Medkova & Cho, 1998; Stahelin et al., 2004; Van Meer et al., 2008). Additionally, phorbol esters are occasionally used as PKC activators while others use the physiological activator DAG despite differences in PKC domain affinity for each activator, potentially confounding results (Ananthanarayanan et al., 2003; Dries et al., 2007; Giorgione et al., 2006; Johnson et al., 2000; Leventis & Grinstein, 2010; Medkova & Cho, 1998; Stahelin et al., 2004). In our study, we propose that PKCε requires binding to importin ⍺ through its NLS motif to tether to the PM to carry out its role in cell migration and invasion.

To investigate the interaction between PKCε and importin ⍺, we first confirmed binding at the predicted NLS sequence through IP of wild-type and NLS-mutant PKCε. Importin ⍺ was shown to bind wild-type PKCε and not the NLS-mutant PKCε validating its binding at the predicted NLS sequence. We then demonstrated that PKCε membrane association was linked to importin ⍺ palmitoylation status. Localization of PKCε to the PM diminished upon inhibition of importin ⍺ palmitoylation. PM localization of both PKCε and importin ⍺ was rescued through the use of importin ⍺-CaaX constructs which target the PM regardless of palmitoylation status (**Fig. 4**). These data demonstrate that PKCε requires binding to palmitoylated importin ⍺ to localize to the PM.

Finally, to link PKCε localization to the phenotypic changes associated with regulating importin ⍺ palmitoylation status, we treated cells with DCP-LA and Wnt-C59 in combination and independently and assessed cell motility. DCP-LA activates PKCε by binding to its C2 domain without enhancing translocation to the PM (Kanno et al., 2015). In the mesenchymal metastatic breast cell line Py230, treatment with DCP-LA did not affect cell motility regardless of Wnt-C59 treatment (**Fig. 5**). This inability to activate PKCε could be indicative of PKCε being maximally active in these tumor cells. We repeated the experiment in the benign breast epithelial cell line, EpH4, which showed that activation of PKCε with DCP-LA treatment increased cell motility. This observed increase was attenuated through additional Wnt-C59 treatment suggesting that preventing importin ⍺ and therefore PKCε from reaching the PM can prevent cell motility. These findings can also explain the observed increase of PKCε at the PM and the increased motility when importin ⍺-CaaX transfected cells were treated with Wnt-C59 (**Fig. 3, 4)**. With PKCε being maximally activated in these tumor cells, an increase in PKCε at the membrane could potentially result in an increase in motility, as observed. It is possible that the population of membrane associated PKCε is increased in the importin ⍺-CaaX cells because CaaX-modified proteins are prenylated which is a permanent modification; this permanent modification may trap PKCε at the membrane as there would be less recycling of importin ⍺ back into the cytoplasm. Furthermore, the increased motility observed with Wnt-C59 treatment in importin ⍺-CaaX cells is an opposite response to the same inhibitor in importin ⍺-WT cells or untransfected cells. This result argues against the hypothesis that the decrease in motility in importin ⍺-WT or untransfected cells treated with Wnt-C59 is a result of downstream effects of Wnt inhibition, such as inhibited non-canonical Wnt signaling or reduced DAG production and consequent lowered PKCε activation. If those downstream effects drove the change in motility, Wnt-C59 should suppress motility regardless of importin ⍺ status, therefore the divergent response in importin ⍺-CaaX cells points to an importin ⍺-dependent mechanism rather than a general consequence of Wnt inhibition.

Our findings describe a novel interaction between importin ⍺ and PKCε and introduce PKCε as a major driving factor for individual and population breast cancer cell motility. We demonstrate a breast cancer cell-specific reliance on PKCε for motility making it an attractive drug target for future studies.

## Acknowledgements

We acknowledge Zainab Ahmed for her contributions to data collection during her rotation in our lab. We thank Dongyan Tan, Camila Dos Santos, and John Haley for their insights and suggestions on experimental approach. We thank the Stony Brook University Department of Pharmacological Sciences (Organization ID: stony-brook-university-pharmsci) for the BioRender license to generate all schematics in this manuscript.

## Author contributions

**M. Kathryn Malone**: Data curation; Formal analysis; Investigation; Methodology; Writing – original draft; Writing – review and editing. **Christopher W. Brownlee:** Conceptualization; Resources; Supervision; Funding acquisition; Project administration; Writing – review and editing.

## METHODS

### Cell Lines, Culture Conditions, and Drug Treatments

MDA-MB-231 (HTB-26, ATCC) and EpH4 (CRL-3063, ATCC) cells were maintained in DMEM high glucose (SH30243.01, Cytiva) supplemented with 10% heat-inactivated fetal bovine serum (FBS) (35-087-CV, Corning). Py230 (CRL-3279, ATCC) cells were maintained in F-12K medium (10-025-CV, Corning) supplemented with 0.1% MITO+ Serum Extender (355006, Corning) and 5% FBS. All media were additionally supplemented with 1% antibiotic-antimycotic (15240062, ThermoFisher). All cell lines were cultured at 37°C in a 90% humidified incubator with 5% CO_2_. During drug treatments, cells were treated with DMSO (40470004, Spectrum Chemical), 10µM Wnt-C59 (S7037, Selleck Chem), 50µM palmostatin (178501, Sigma-Aldrich), 40µM importazole (S8446, Selleck Chem), 25µM ivermectin (J6277.03, ThermoFisher), 100µM 2-bromopalmitate (21604, Sigma-Aldrich), or 500nM DCP-LA (HY-108599A, MedChemExpress) at 37°C for 24 hours

### Imaging

All cell imaging was performed using an EVOS M7000 epifluorescence and transmitted light microscope. Live cell imaging was performed at 37°C with 70-90% humidity and 5% CO_2_ using an EVOS Onstage Incubator. Images were taken with an Olympus UPLFLN 4x/numerical aperture 0.13, Olympus UPLFLN 20x/numerical aperture 0.50, or Olympus UPLFLN 40x/numerical aperture 0.75 objective. Images taken as z-stacks were processed using 3D deconvolution from Celleste Image Analysis Software.

### Wound Closure Assay

Cells were plated to form a confluent monolayer and were scratched with a 200 μL pipette tip to create a cell-free area. Cells were then incubated in complete media in the presence or absence of drugs and images were taken every 6 hours for 24 hours post scratch. Wound closure percentage was calculated as the difference between the initial scratch area and remaining cell-free area divided by the initial scratch area.

### Transwell Invasion Assay

Cell invasion was assessed using Matrigel (356234, Corning)-coated Transwell (3470, Corning) inserts. Cells were seeded at a concentration of 5×10^4^ cells/mL in serum-free media into the upper chamber of the Transwell inserts. Complete media was added to the lower chamber as a chemoattractant. Cells were incubated for 24 hours in the presence or absence of drugs. At 24 hours, cells were removed from the inner membrane and those adhered to the outer membrane were methanol fixed and stained with crystal violet. Whole membranes were imaged and relative area was calculated as a ratio of cell-positive area to total membrane area.

### Cell Tracking

Cell tracking was performed as previously described (Pijuan et al. 2019). Cells were plated immediately following transfection. 24 hours post transfection, drug treatments were administered and imaging began. Images were taken at 20-minute intervals for 20 hours.

### Intracellular Localization and Nuclear Localization Signal Sequence prediction

Genes identified by Simpson et al. to impair cell migration upon siRNA knockdown were further screened using UniProt GO identifiers (Ashburner et al. 2000, Thomas 2017) for their localization at the plasma membrane and the cytosol. The three nuclear localization signal predictors NucPred (Brameier et al. 2007), NLStradamus (Nguyen et al. 2009), and cNLSmapper (Kosugi et al. 2009) were then used to predict the presence and location of potential NLS sequences in each gene.

### Cell Lysis, Co-Immunoprecipitation, and Western Blot Analysis

Cells were grown on 10cm dishes and were collected at 90% confluence by scraping following an ice-cold PBS wash. The cell suspension was spun at 500rpm for 5 minutes and the pellet was resuspended in non-denaturing lysis buffer (20mM Tris HCl pH8, 137mM NaCl, 1% Triton X-100, 2mM EDTA) supplemented with 10mg/mL of leupeptin, pepstatin, and chymostatin protease inhibitors. The lysate was then rocked for 1 hour at 4°C and spun at 12,000rpm for 20 minutes at 4°C. The supernatant was then removed and diluted with 2x Laemmli buffer and boiled at 100°C for 5 minutes and stored at - 20°C until use. Co-immunoprecipitation was performed using ThermoFisher Pierce anti-HA magnetic beads (88836, Thermo Scientific) following the recommended protocol. Protein products were analyzed via western blot by running through SDS-PAGE and transferring to nitrocellulose membranes and blotting for target proteins. Rabbit polyclonal anti-KPNA2 (10819-1-AP, Proteintech), rabbit polyclonal anti-HA tag (51064-2-AP, Proteintech), Rabbit polyclonal anti-GAPDH (10494-1-AP, Proteintech), and goat anti-rabbit IgG AF-700 (A21038, Invitrogen) antibodies were used.

### DNA Mutagenesis, Plasmids, and Nucleofection

The plasmids UBC-hKPNA2-HA-mCherry and UBC-hKPNA2-HA-mCherry-CaaX were designed with and purchased from VectorBuilder. PKC epsilon WT (PKCε-HA) was a gift from Bernard Weinstein (Addgene plasmid # 21240). This plasmid was used for WT PKCε experiments and was used as the backbone for the PKCε-HA-ΔNLS mutant. The PKCε-HA-ΔNLS mutant was generated using the Q5 site-directed mutagenesis kit (E0554S, New England Biolabs) with primer sets F: 5’-aaagcgCTCGCTGCTGGTGCTG-3’ and R: 5’-gccgccTTGGCCACTGTTGGTG-3’ with 2% DMSO following the recommended protocol. Py230 cells were transfected via nucleofection using Lonza SG cell line 4-D nucleofector kit with the pulse code EN-150, both of which were optimized for this cell line. 4×10^5^ cells and 1µg plasmid were used with 16-well Nucleocuvette® Strips; 2×10^6^ cells and 5µg plasmid were used with 100µL Nucleocuvette® Vessels. MDA-MB-231 cells were transfected via nucleofection using the recommended Lonza SE cell line 4-D nucleofector kit with the pulse code CH-125. 1×10^5^ cells and 400ng plasmid were used with 16-well Nucleocuvette® Strips; 5×10^5^ cells and 2µg plasmid were used with 100µL Nucleocuvette® Vessels. Both cell lines were plated immediately following transfection and incubated for 24 hours prior to use.

### Immunostaining and Mitotic Analysis

Cells were plated on fibronectin-coated coverslips and were incubated for 24 hours. Cells were then fixed with 4% PFA, permeabilized with PBS+0.2% Triton X-100 (9036-19-5, Sigma-Aldrich), and blocked with 3% Bovine Serum Albumin (BSA) in PBS+0.2% Triton X-100. Antibodies were diluted in PBS+0.2% Triton X-100 at the following concentrations: rabbit polyclonal anti-phospho-histone H3 (pSer^10^) 1:1000 (H0412, Sigma-Aldrich), mouse monoclonal anti-HA tag 1:1000 (SAB2702196, Sigma-Aldrich), rabbit polyclonal anti-PKCε 1:1000 (20877-1-AP, Proteintech), donkey anti-rabbit IgG AF-488 1:1000 (6440-30, Southern Biotech), donkey anti-mouse IgG AF-488 1:1000 (6411-30, Southern Biotech), donkey anti-mouse IgG AF-568 1:1000 (A10037, Invitrogen). The coverslips were then mounted onto slides with ProLong Diamond Antifade Mountant (P36961, ThermoFisher). Mitotic analyses were performed on cells following fixation and immunostaining with anti-phospho-histone H3 antibody. Mitotic ratio was calculated as the ratio of phospho-histone H3 positive cells to total cells counted.

### Plasma membrane to cytoplasm ratio analysis

Deconvolved images were analyzed using ImageJ (v2.16.0). A 120×10 pixel straight line was used to measure the integrated density of HA tag along the PM and in the cytoplasm of each cell.

### Quantification and statistical analysis

All statistical analyses were performed using GraphPad Prism (v10.6.1). Statistical details can be found in the figure legends. Results are expressed as mean ±SEM or as median with quartiles, ns = not significant, *P<0.05, **P<0.01, ***P<0.001, ****P<0.0001.

